# Expression of the Eight GABA_A_ Receptor α Subunits in the Developing Zebrafish Central Nervous System

**DOI:** 10.1101/244590

**Authors:** Bryan Monesson-Olson, Jon J. McClain, Abigail E. Case, Hanna E. Dorman, Daniel R. Turkewitz, Aaron B. Steiner, Gerald B. Downes

## Abstract

GABA is a robust regulator of both developing and mature neural networks. It exerts many of its effects through GABA_A_ receptors, which are heteropentamers assembled from a large array of subunits encoded by distinct genes. In mammals, there are 19 different GABA_A_ subunit types, which are divided into the α, β, γ, δ, ε, π, θ and ρ subfamilies. The immense diversity of GABA_A_ receptors is not fully understood. However, it is known that specific isoforms, with their distinct biophysical properties and expression profiles, tune responses to GABA. Although larval zebrafish are well-established as a model system for neural circuit analysis, little is known about GABA_A_ receptors diversity and expression in this system. Here, using database analysis, we show that the zebrafish genome contains at least 23 subunits. All but the mammalian θ and ε subunits have at least one zebrafish ortholog, while five mammalian GABA_A_ subunits have two zebrafish orthologs. Zebrafish contain one subunit, β4, which does not have a clear mammalian ortholog. Similar to mammalian GABA_A_ receptors, the zebrafish α subfamily is the largest and most diverse of the subfamilies. In zebrafish there are eight α subunits, and RNA *in situ* hybridization across early zebrafish development revealed that they demonstrate distinct patterns of expression in the brain, spinal cord, and retina. Some subunits were very broadly distributed, whereas others were restricted to small populations of cells. Subunit-specific expression patterns were similar between zebrafish, frogs and rodents, which suggests that the roles of different GABA_A_ receptor isoforms are largely conserved among vertebrates. This study provides a platform to examine isoform specific roles of GABA_A_ receptors within zebrafish neural circuits and it highlights the potential of this system to better understand the remarkable heterogeneity of GABA_A_ receptors.

## INTRODUCTION

Neural networks throughout the central nervous system rely upon a diversity of neurotransmitter systems for both their initial formation and mature function. GABA is the major inhibitory neurotransmitter throughout most of the mature nervous system [1]. It exerts its effects through its receptors, which are divided into two classes. GABA_A_ receptors are ligand-gated chloride channels responsible for most of the rapid effects of GABA, while GABA_B_ receptors are heterotrimeric G-protein coupled receptors. In mammalian systems, GABA_A_ receptors demonstrate incredible subtype diversity and are targets for classes of clinically important drugs, such as benzodiazepines and barbituates [2, 3]. GABA_A_ receptors are heteropentamers composed of various combinations of 19 different subunits: α1–6, β1-3, γ1-3, δ, ε, π, θ and ρ1-3 (previously referred to as GABA_c_ receptors) [4, 5]. Each subunit is encoded by a discrete gene that is spatially and developmentally regulated to generate distinct, but often overlapping, expression patterns [6–9]. Alternative splicing and RNA editing of some subunits further increases the number of subtypes available [10]. Although this extensive receptor heterogeneity is not fully understood, some subunits confer distinct biophysical and pharmacological properties, interact with specific cytoplasmic proteins, and localize to specific subcellular domains [11–13]. Ultimately, this receptor diversity provides a capacity to tailor responses to GABA within neural circuits.

The α subunits play a prominent role in GABA_A_ receptor function. They are thought to be essential components of ‘classic’ GABA_A_ receptors, which exclude the pharmacologically distinct receptors composed from ρ subunits [3]. Exogenous expression studies and analysis of native receptors indicate that each receptor contains two α subunits which, along with β subunits, form the GABA binding site. α subunits can also dictate isoform-selective pharmacology. For example, receptors containing α1, α2, α3 and α5 subunits confer benzodiazepine-sensitivity, whereas receptors that contain α4 and α6 subunits do not bind to clinically used benzodiazepines [2]. In addition, α subunits regulate subcellular localization. GABA receptors that contain α1, α2, and α3 demonstrate a synaptic localization and mediate transient or phasic inhibition, while receptors that contain α4, α5 and α6 are principally extrasynaptic and mediate tonic inhibition [14, 15]. Through both transient and tonic inhibition, these receptors mediate the majority of inhibition in neural circuits throughout the vertebrate brain.

Developing zebrafish have several features that make them well-suited to study the formation and function of neural circuits [16, 17]. First, zebrafish embryos and larvae develop external to the mother and are optically transparent. These features make the central nervous system easily accessible throughout development and particularly amenable to optical physiological approaches, such as optogenetics or calcium imaging. Second, the central nervous system contains fewer cells compared to mammalian preparations, yet many cell types and mechanisms are conserved among vertebrates. Lastly, larval zebrafish develop rapidly, therefore many sensory and motor systems are present and functional within five days post-fertilization. Combined, these features have helped establish developing zebrafish as a prominent system to examine neural circuit formation and function.

GABA_A_ receptors are expressed in larval zebrafish and essential for normal nervous system function. For example, bath application of pentylenetetrazole, a convulsant and noncompetitive GABA_A_ receptor antagonist, induces hyperactive behavior that can serve as a model of seizures [18]. Similarly, insecticides known to target GABA_A_ receptors generate hyperactive behavior in larvae [19]. Pharmacological blockade of GABA_A_ receptors has also been shown to regulate the activity of specific cells, including retinal bipolar cells [20], optic tectum neurons [21], Mitral cells in the olfactory bulb [22], spinal cord CoPA interneurons and motoneurons [23], and the Mauthner cells, a pair of well-studied reticulospinal neurons found in the amphibian and teleost hindbrain [24]. In these studies, the GABA_A_ receptor isoforms were not identified. However, patch-clamp recordings of the Mauthner cells showed three distinct GABA_A_ kinetic profiles, which were proposed to be caused by different receptor isoforms [25].

There is limited information about the heterogeneity of GABA_A_ receptors in zebrafish. The most extensive study to date identified 23 GABA_A_ receptor subunits and examined their expression broadly in adult zebrafish brain via RT-qPCR [26]. Despite playing prominent roles in developing animals, little is known about the expression of GABA_A_ receptor subunits in zebrafish embryos and larvae. α1 has been shown to be expressed at 50 and 72 hours post-fertilization (hpf), γ2 at 50 hpf, and α2a (then named α2) from ~14 – 96 hpf. [27–29]. Unpublished α6a expression data has also been deposited in the ZFIN database [30]. The expression patterns of other GABA_A_ receptor subunits has not been reported.

In this study we performed phylogenetic analysis, which indicates that zebrafish contain at least 23 different GABA_A_ receptor subunits. Although we observed some differences between the zebrafish and mammalian GABA_A_ receptor subunit gene families, zebrafish contain orthologs for most of the GABA_A_ receptor subunits found in mammals. To examine where and when these GABA_A_ receptors are expressed in the zebrafish nervous system, we performed whole-mount RNA *in situ* hybridization. We present the embryonic and larval expression of the eight identified α subunit-encoding genes: *gabra1*, *gabra2a*, *gabra2b*, *gabra3*, *gabra4*, *gabra5*, *gabra6a* and *gabra6b*. These data show that each subunit has a distinct expression pattern, broadly similar to the reported expression of amphibian and rodent orthologs. Combined, these results suggest that GABA_A_ isoform-specific roles are conserved among vertebrates, and they establish a foundation to use the zebrafish system to better understand how GABA_A_ receptor heterogeneity mediates neural circuit formation and function.

## MATERIALS AND METHODS

### Phylogenetic analysis

Homologous gene queries were performed using the National Center for Biotechnology Information tBLASTn search tool. Protein sequences for each of the 19 mouse GABA_A_ receptor subunits were used to search the zebrafish nucleotide collection. Matching zebrafish sequences were evaluated to determine if they were splice variants from the same gene or generated from different genes. When splice variants were identified, only the longest variant was used for subsequent analysis. 19 mouse and 23 zebrafish protein sequences were used to assemble a phylogenetic tree. The genes and Genbank accession numbers for the mouse sequences are: *Gabra1*- NP_034380, *Gabra2*- NP_032092, *Gabra3*- NP_032093, *Gabra4*- NP_034381, *Gabra5*- NP_795916*, Gabra6*- NP_00109311, *Gabrb1*- NP_032095, *Gabrb2*- NP_032096, *Gabrb3*- NP_032097, *Gabrd*- NP_032098, *Gabre*- NP_059065, *Gabrg1*- NP_034382, Gabrg2-NP_032099, *Gabrg3*- NP_032100, *Gaprp*- NP_666129, *Gabrq*- NP_065234, *Gabrr1*-NP_032101, *Gabrr2*- NP_032102, and *Gabrr3*-NP_001074659. The Genbank accession numbers for the zebrafish sequences are: *gabra1*- NM_001077326, *gabra2a*- XM_009307207, gabra2b- XM_017359049, *gabra3*- XM_009295708, *gabra4*- NM_001017822, *gabra5*-XM_001339475, *gabra6a*- XM_005173112, *gabra6b*- XM_002667357, *gabrb1*- XM_002664133, *gabrb2*- NM_001024387, *gabrb3*- XM_005166079, *gabrb4*- XM_005173874, *gabrd*-XM_695007, *gabrg1*- XM009307208, *gabrg2*- NM_001256250, *gabrg3*- XM_009302568, *gabrp*-XM_005173293, *gabrr1*- NM_001025553, *gabrr2a*- NM_001045376, *gabrr2b*- XM_009294512, *gabrr3a*- NM_001128760, *gabrr3b*- XM_009297450, *gabrz*-XM_005156247. To construct the tree, a ClustalW alignment of amino acid sequences was performed using the Geneious software package [31]. An unrooted PHYML consensus tree was then generated using 100 bootstrap replicates [32].

### Fish maintenance and breeding

Zebrafish were raised and maintained using established husbandry procedures. Embryos were kept at 28.5 °C in E3 media and staged according to morphological criteria [33]. All experiments were performed using Tuebingen (Tu) or tub longfin (TLF) wild type embryos. From 0–24 hpf, embryos were grown in 0.01% Methylene Blue in E3 medium. At 24 hpf the solution was changed to 0.0045% Phenylthiourea (PTU) in E3 to inhibit pigment formation. This solution was changed every 24 hours until the fish were sacrificed. All animal protocols were approved by the University of Massachusetts Institutional Animal Care and Use Committees (IACUC).

### Whole-mount in situ hybridization

Antisense digoxigenin probes were generated against *gabra1*, *gabra2a*, *gabra2b*, *gabra3*, *gabra4*, *gabra5*, *gabra6a* and *gabra6b* (S1 Table). *In situ* probe synthesis utilized the digoxigenin RNA labeling kit with SP6 or T7 RNA polymerases (Roche Diagnostics, Mannheim, Germany). Whole-mount, colorimetric *in situ* hybridization was performed using established protocols [34] and examined using a compound microscope (Zeiss, Thornwood, NY) attached to a digital camera (Zeiss, Thornwood, NY). Cross sections were generated by imbedding *in situ* hybridization stained embryos in 1.5% agar, 5% sucrose. The blocks were submerged in a 30% sucrose solution overnight then cut into 20mm thick sections using a cryostat (Leica, Buffalo Grove, IL). Representative larvae at 24, 48, and 96 hpf were mounted in either 100% glycerol or dehydrated through a methanol series, equilibrated in a 2:1 benzylbenzoate/benzylalchol solution and mounted in a 10:1 Canada balsam/methyl saliylate misture. Images were captured using a Zeiss Discovery v12 stereomicroscope with a 1.5x objective or a Zeiss Axioskop Microscope with a 10x objective and an AxioCam MRc camera. All images were processed using Adobe Photoshop, in which multiple focal planes were merged to produce single representative images. The identify of neuroanatomical structures was determined using zebrafish brain atlases [35–37].

### Reverse-Transcriptase PCR

RT-PCR was used to analyze whether the *gabra3* and *gabra3*-like sequence were portions of the same transcript. Primers designed against *gabra3*-like (Primer 1 5’ GGACGGCGGATGATGAGAAA-3’) and *gabra3* (Primer 2 5’-CACGACCGTCCTGACTA-3’, Primer 3 5’-GTGGAGTAGATGTGGTGGGC-3’) were used to amplify cDNA from wild-type zebrafish larvae. RNA was extracted from 48 hpf zebrafish larvae using the RNAeasy kit (Qiagen, Venlo, Netherlands) and reverse transcribed using the Accuscript RT-PCR system (Stratagene). The PCR products were sequenced to confirm that they are portions from the same transcript.

## RESULTS

### The zebrafish GABA_A_ receptor subunit gene family is similar in size and diversity to the mammalian GABA_A_ receptor gene family

19 GABA_A_ receptor subunit genes have been identified in humans, mice, and rats [4, 5]. To establish the number and organization of zebrafish GABA_A_ receptor subunits, we used mouse GABA_A_ receptor subunit amino acid sequences to query zebrafish genome databases. We identified 23 zebrafish GABA_A_ receptor subunits, each encoded by a distinct gene. Frequently splice variants were observed. In these cases, the principle splice isoform, as indicated by the databases, was selected for analysis. 12 of the 19 mouse GABA_A_ receptor subunits were found to have a single ortholog in zebrafish (Fig 1). Amino acid identities between mouse and zebrafish orthologs ranged from 53.1 to 86.3%. Five additional mouse subunits, α2, α6, ρ2, ρ3, and π, each exhibit similarity with two subunits in zebrafish: *gabra2a* and *gabra2b*, *gabra6a* and *gabra6b*, *gabrr2a* and *gabrr2b*, *gabrr3a* and *gabrr3b*, *gabrp* and *gabrz*, respectively. The relatively high percentage of amino acid identity of these zebrafish subunits with each other suggests that the zebrafish paralogs are duplicated genes. Duplicated genes are often observed in zebrafish, and they are thought to be due to a whole genome duplication within the teleost lineage [38, 39]. It was reported previously that there is a *gabra3* and an α3-like gene. While the α3-like gene has been localized to chromosome 21, the genomic location of the α3-like gene is not known. Our database analysis suggested that these sequences could be non-overlapping portions of the same gene, therefore RT-PCR was performed using primers targeting α3-like and *gabra3* sequences (S1 Fig). Transcripts were amplified that spanned the region between *gabra3* and α3-like sequences, indicating that these are different portions of the same gene and that there is only one α3 encoding gene, *gabra3*.

**Fig 1.**
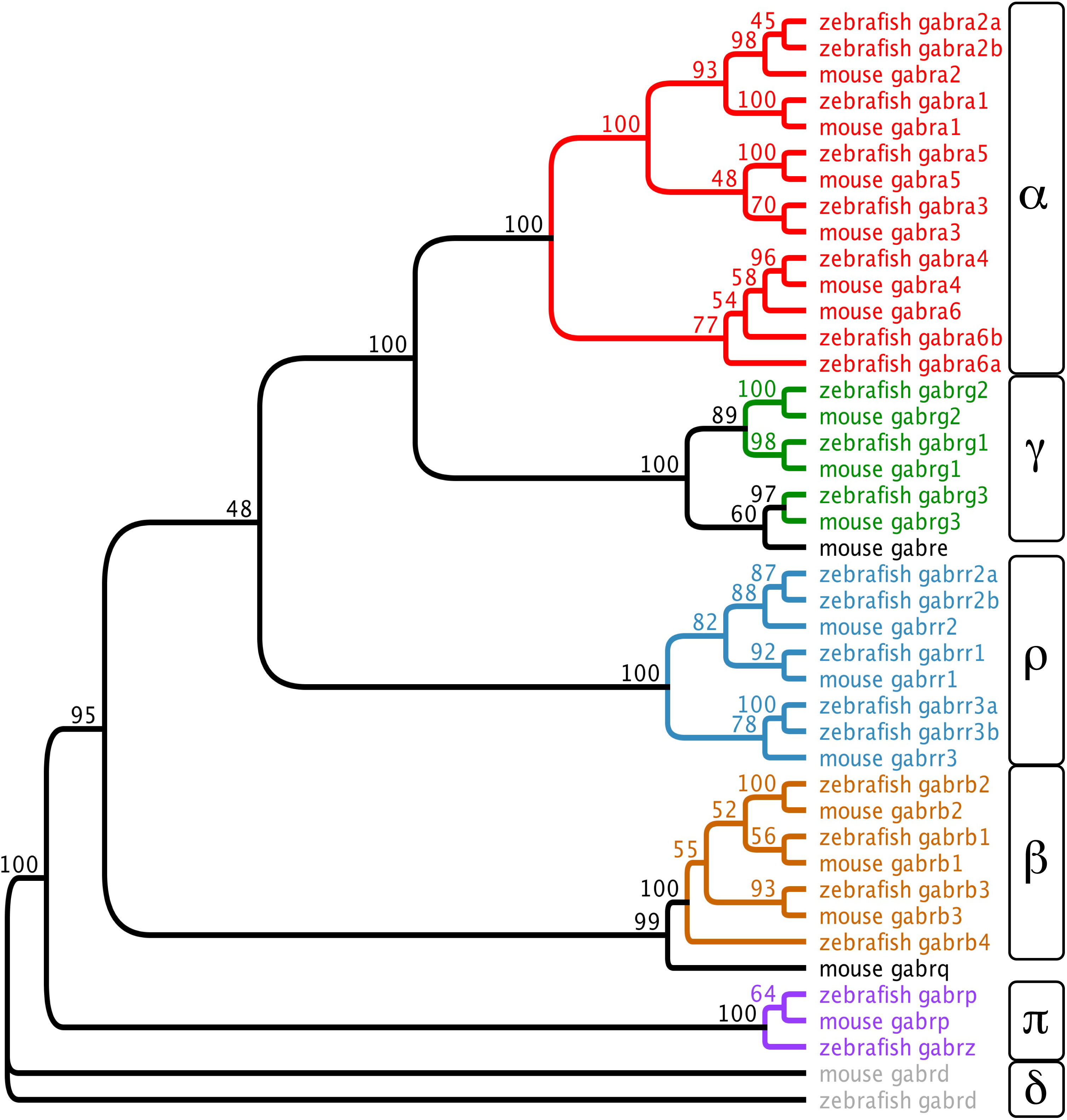
Phylogenetic analysis shows that the zebrafish GABA_A_ subunit gene family is similar in size, diversity, and organization to the mouse GABA_A_ subunit gene family. Amino acid sequence alignments were used to generate a consensus tree using 100 bootstrap replicates. The genes that encode the GABA_A_ subunits are shown at the tip of each branch and bootstrap proportions are shown at the branch points. The zebrafish and mouse GABA_A_ subunit sequences showed high amino acid identity and grouped into the α, β, γ, δ, π, and ρ subfamilies. Most mouse GABA_A_ subunits have a single zebrafish ortholog, while six mouse GABA_A_ subunits have two zebrafish orthologs, likely due to gene duplication.

There are two mouse GABA_A_ receptor subunits for which zebrafish orthologous are less clear or have yet to be identified, the θ and ε subunits. The θ subunit, encoded by the gene *Gabrq*, exhibits only 31.3% amino acid identity with zebrafish β4, encoded by *gabrb4*, which is the greatest percentage identity of this subunit with any zebrafish sequence. There are three β subunits in mice compared to the four β subunits in zebrafish so it is possible that, despite the low amino acid sequence identity, zebrafish β4 is orthologous to the mouse θ subunit. No clear zebrafish ortholog has yet to be identified for the mouse ε subunit, which is encoded by the *Gabre* gene.

### α subunits demonstrate distinct expression patterns across early zebrafish development

Determining the temporal and spatial expression of GABA_A_ subunits is instrumental to identify isoform-specific roles in zebrafish neural circuit development and function. Therefore, given the large size of the α subunit subfamily and their prominent role in GABA_A_ receptor function, we examined the expression of the eight α subunit encoding genes: *gabra1*, *gabra2a*, *gabra2b*, *gabra3*, *gabra4*, *gabra6a* and *gabra6b*. Using whole-mount RNA *in situ* hybridization, we determined their expression patterns at 24, 48, and 96 hpf. These time points span much of larval zebrafish locomotor network development [40].

Expression of *gabra1*, *gabra2a*, *gabra2b*, *gabra3*, *gabra4* and *gabra5* was detected at 24 hpf (Fig 2). Neither *gabra6a* or *gabra6b* were detected at this developmental stage. *gabra2a* and *gabra2b* were detected broadly and did not appear to be spatially restricted (Fig 2C-F). To distinguish whether these gene transcripts were widespread or background due to the probe, a second probe was generated for each gene (S1 Table). These probes yielded widespread staining, very similar to the initial probes used, which suggests that *gabra2a* and *gabra2b* are widely distributed. Consistent with these findings, a previous study reported that *gabra2a* exhibits a diffuse, broad pattern of expression in larval zebrafish [28].

**Fig 2.**
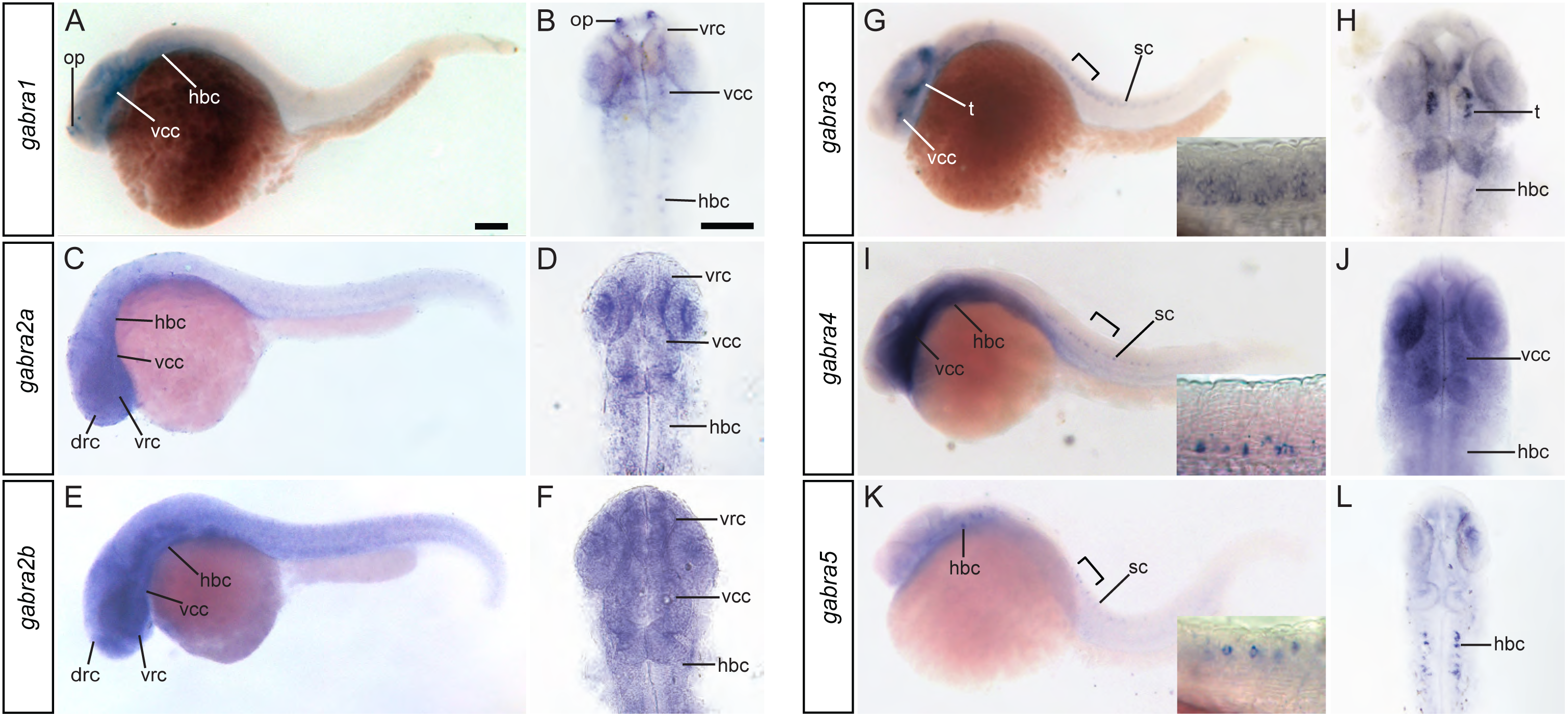
Expression of GABA_A_ α subunits at 24 hpf. *gabra1* (A, B), *gabra2a* (C, D), *gabra2b* (E, F), *gabra3* (G, H), *gabra4* (I, J), and *gabra5* (K, L) were detected at this time point. Whole mount lateral (A, C, E, G, I, K) views are shown along with dorsal views of the head (B, D, F, H, J, L).The scale bar (A) is 0.1 mm. The brackets in G, I, and K indicates the regions shown at higher magnification in the corresponding insets. Abbreviations: drc, dorsal rostral cluster; hbc, hindbrain cluster; op, olfactory placode; SC, spinal cord; t, tegmentum; vcc, ventral caudal cluster; vrc, ventral rostral cluster.

*gabra1*, *gabra3*, *gabra4*, and *gabra5* were each expressed in discrete cells in the brain or spinal cord at 24 hpf. *gabra1* was detected in the olfactory placodes, the ventral rostral cluster, the ventral caudal cluster and small clusters of cells within each rhombomere of the hindbrain (Fig 2A, B). Although expressed in discrete cells, *gabra3* was widely expressed, and found within the tegmentum, hindbrain, and ventral and intermediate domains in the spinal cord (Fig 2G, H). *gabra4* was expressed in a diffuse manner throughout the brain but specifically within a select population of ventral spinal cord cells (Fig 2I, J). The location and distribution of these cells suggests that *gabra4* is expressed selectively within Kolmer-Agduhr (KA) neurons. Lastly, *gabra5* is prominently expressed in small groups of cells in each rhombomere of the hindbrain and in an intermediate domain of the spinal cord (Fig 2K, L). The location and distribution of these spinal cord cells suggests that they are CoPA neurons.

The expression of all GABA_A_ receptor subunit genes, except for *gabra6a*, was observed at 48 hpf (Fig 3). Similar to the pattern observed at 24 hpf, the *gabra2* paralogs were again detected broadly, with little spatial restriction (Fig 3C-F). Similar to its expression at 24hpf, *gabra1* is prominently expressed in the olfactory bulbs, the pallium and discrete clusters of cells in medial portions of the medulla (Fig 3A, B). *gabra3* was observed in relatively small clusters of cells in the pallium, thalamus, and the medulla (Fig 3G, H). *gabra4* transcripts were identified in distinct cells in the subpallium and lateral potions of the medulla. *gabra5* transcripts were detected most prominently in the medulla. The location and expression this subunit suggests it exhibits robust expression in the Mauthner cells, therefore α5 containing receptors are likely some of the GABA_A_ isoforms identified previously via electrophysiology [25]. *gabra6b* is expressed prominently in the olfactory bulbs.

**Fig 3.**
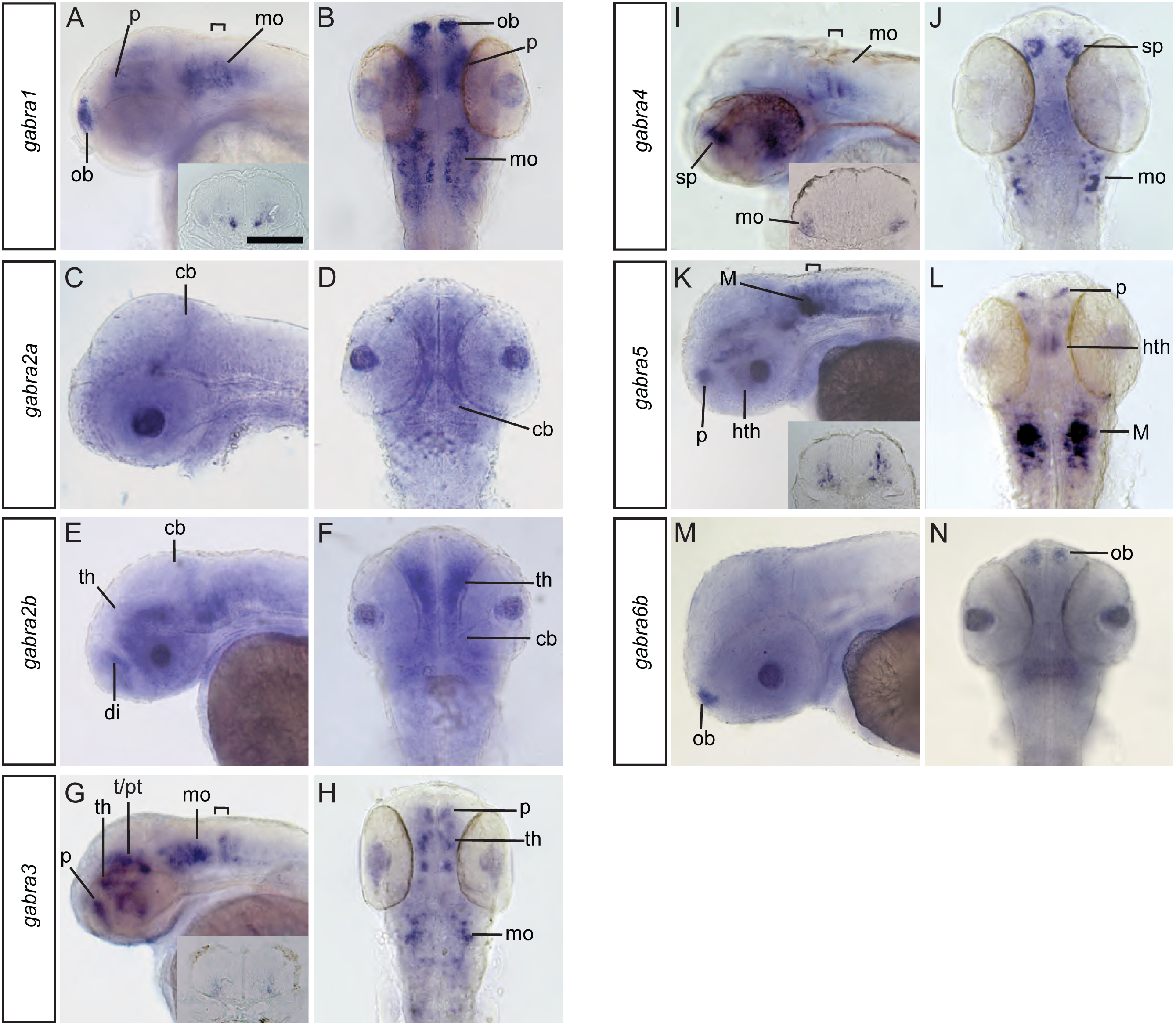
Expression of the GABA_A_ α subunits in the brain at 48hpf. *gabra1* (A, B), *gabra2a* (C, D), *gabra2b* (E, F) *gabra3* (G, H) *gabra4* ( I, J) *gabra5* (K, L) and *gabra6b* (M, N) were detected at this time point. Lateral (A, C, E, G, I, K, M) and dorsal (B, D, F, H, J, L, N) views are shown. The scale bar (A) is 0.1 mm. Brackets in A, G, I, and K indicate the region shown in cross sections within the insets. Abbreviations: cb, cerebellum; di, diencephalon; hth, hypothalamus; M, Mauthner cell; mo, medulla oblongata; ob, olfactory bulb; p, pallium; sp, subpallium; t, tegmentum; th, thalamus.

Although *gabra1*, *gabra3*, *gabra4*, and *gabra5* transcripts were all detected in the medulla, they do not appear to be in the same cells. Instead, an intriguing medial to lateral organization was observed. *gabra1* is expressed strongly in the most medial cells, *gabra3* and *gabra5* transcripts were found in an intermediate domain, and *gabra4* was observed laterally (compare insets in Fig A, G, I, K).

Transcripts for all α subunits are detected in larval zebrafish at 96 hpf (Fig 4). As with the earlier developmental time points, *gabra2a* and *gabra2b* transcripts were detected broadly, although *gabra2b* appears less diffuse and more discrete compared to at 24 and 48 hpf (Fig 4C-F). *gabra1*, *gabra3*, and *gabra6b* are also expressed more broadly compared to earlier stages. In contrast, *gabra4*, *gabra5* and *gabra6a* transcripts were detected in smaller groups of cells. *gabra4* is expressed in a stripped pattern in the outer nuclear layer of the retina, the posterior tuberculum area, which is a portion of the diencephalon, and the tectum, cerebellum, and medulla (Fig I, J). *gabra5* is expressed in discrete cells in the pallium, hypothalamus, cerebellum and medulla (Fig K, L). Within the medulla, *gabra5* expression in the Mauthner Cells is robust, as it is at earlier developmental stages.

**Fig 4.**
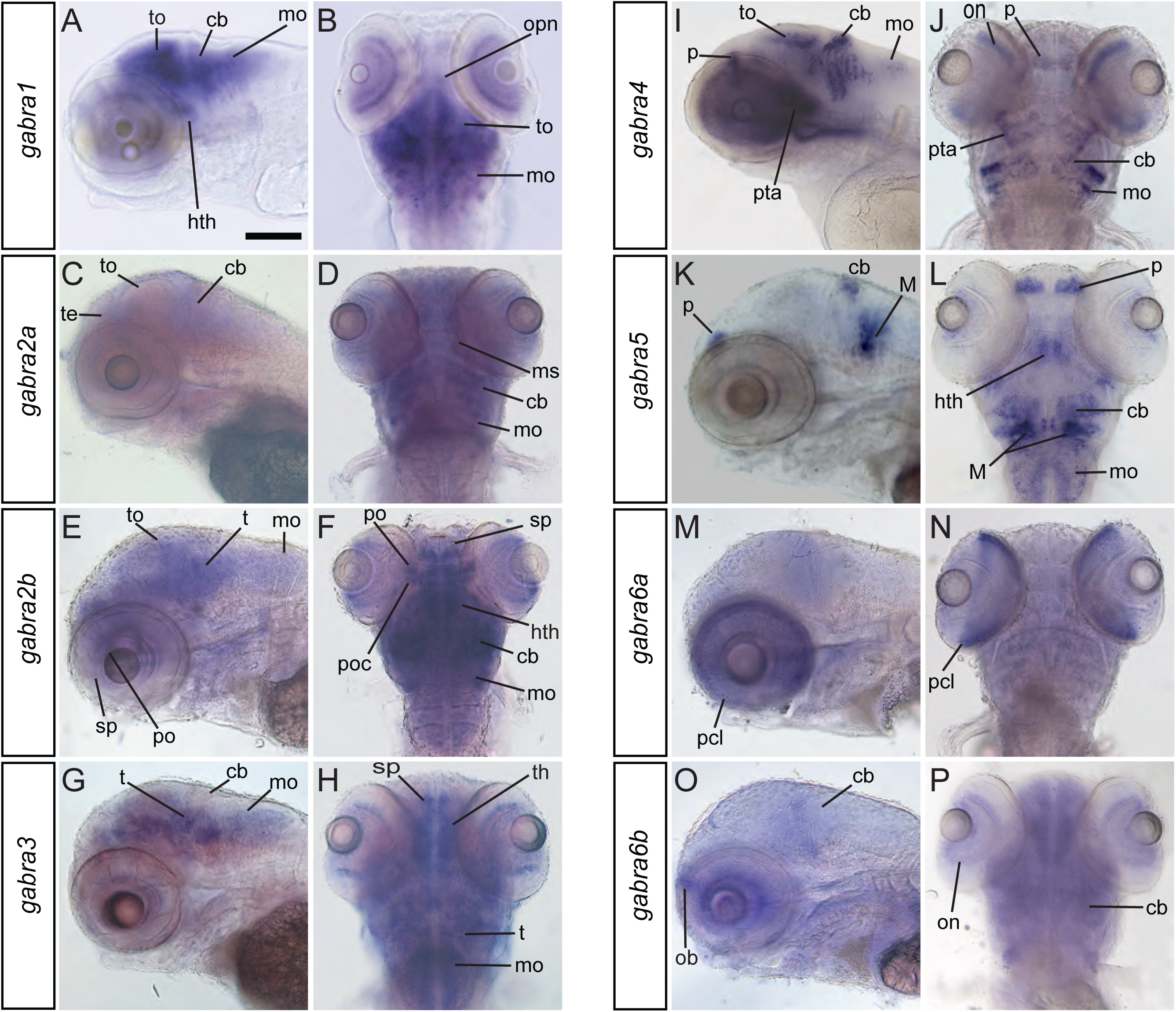
Expression of the GABA_A_ α subunits in the brain at 96hpf. Transcripts encoding all α subunits were detected at this time point. Lateral (A, C, E, G, I, K, M, O) and dorsal (B, D, F, H, J, L, N, P) are shown The scale bar (A) is 0.1 mm. Abbreviations: cb, cerebellum; gcl, ganglion cell layer; hth, hypothalamus; inl, inner nuclear layer;M, mauthner cell; mo, medulla oblongata; ob, olfactory bulb; opn, optic nerve; p, pallium; pcl, photoreceptive cell layer; po, preoptic region; poc, post optic commissure; pta, posterior tubercular area; sp, subpallium; t, tegmentum; te, telencephalon; th, thalamus; to, tectum opticum;

Transcripts for several α subunits are expressed in the retina at 96 hpf, however most are restricted to one or two cell layers (Fig 4). *gabra1* and *gabra6b* are both expressed in the ganglion cell and outer nuclear layers, *gabra2a* and *gabra2b* both seem to be expressed broadly, and *gabra3* and *gabra4* transcripts both are restricted to the outer nuclear layer. *gabra6a* is expressed prominently in the photoreceptor cell layer, and does not demonstrate robust expression outside of the retina.

## DISCUSSION

In this study, we showed that the GABA_A_ receptor gene family exhibits a size and diversity very similar to those found in mammals. We identified sequences for 23 GABA_A_ receptor encoding genes, and phylogenetic analysis indicates that isoform are conserved among vertebrates for the majority of subunits. We determined the expression of the eight zebrafish α subunit encoding genes, the largest and most diverse gene family, and observed that they are expressed in distinct, often overlapping expression patterns. Taken together, these data argue that larval zebrafish can serve as a useful model to investigate the functional roles of GABA_A_ receptor subtypes within developing neural circuits.

A previous study identified 23 GABA_A_ receptor genes in zebrafish. Our data largely confirm their results, except ρ3b, which is encoded by *gabrr3b*, was not reported. A α3 and an α3-like gene were also described, but our analysis indicates that zebrafish likely contain only one α3 gene. Given the amount of zebrafish genome data that is available, it seems unlikely that additional GABA_A_ receptor subunit encoding genes exist. However, similar to mammalian systems, sequence data indicates that several zebrafish GABA_A_ receptor encoding genes are alternatively spliced, which further enhances the already extensive diversity of receptor isoforms.

The evolution of the GABA_A_ receptor gene family has been examined by using sequence data from a wide variety of species, including humans, rodents, canary, chicken, frogs and pufferfish, tunicates, *C. elegans* and *drosophila* [41, 42]. Our results are in line with these studies in finding that mammalian θ and ε subunits are more distinct compared to other subunits, such that their existence outside of the mammalian lineage is unclear [43]. Even among mammalian species, for example comparing humans to rodents, these two subunits are very diverse, which suggests they are evolving at a much faster rate than other subunits [44]. The roles of θ and ε subunits may be unique to mammals or, alternatively, fulfilled in zebrafish and other species using other subtypes. Experiments to investigate the functional ability of zebrafish subtypes to substitute for θ and ε in rodent systems could distinguish between these possibilities.

The α subunits demonstrate a wide range of expression patterns across early zebrafish development (Table 1). On one end of the spectrum are *gabra2a* and *gabra2b*, which appeared to be expressed very broadly at each of the three time points examined. At the other end of the spectrum is *gabra6a*, which we detected only at 96 hpf and mostly in the photoreceptor cell layer of the retina. The other subunits fall between these extremes, with expression in discrete cells in various brain and spinal cord regions. When GABA_A_ receptor α subtypes are expressed in the same region, they often occupy different domains. For example, *gabra1*, *gabra3*, *gabra4*, and *gabra5* are all expressed in the medulla, but are organized in medial to lateral stripes. The zebrafish medulla has a structural and functional organization in which neurons of shared neurotransmitter, phenotype, age, morphology and functional properties are arranged into stripes [45, 46]. Another example of different GABA_A_ α subtypes occupying different domains within the same region is found in the spinal cord. *gabra4* transcripts were found in ventral cells, likely KA neurons, *gabra5*-expressing cells, likely CoPA neurons, were observed more dorsally, and *gabra3* transcripts were detected more broadly in both ventral and more dorsal domains. It is not yet clear how the expression of different GABA_A_ subtypes correspond to the organizational stripes of the hindbrain or cell types in the spinal cord, but it seems likely that some subtypes demonstrate cell-type specific expression and confer distinct responses to GABA. Conclusive cell-type identification will require colabeling experiments that examine GABA_A_ subunit expression along with specific markers or techniques that determine neuronal identity by revealing cell morphology [47].

**Table 1.**
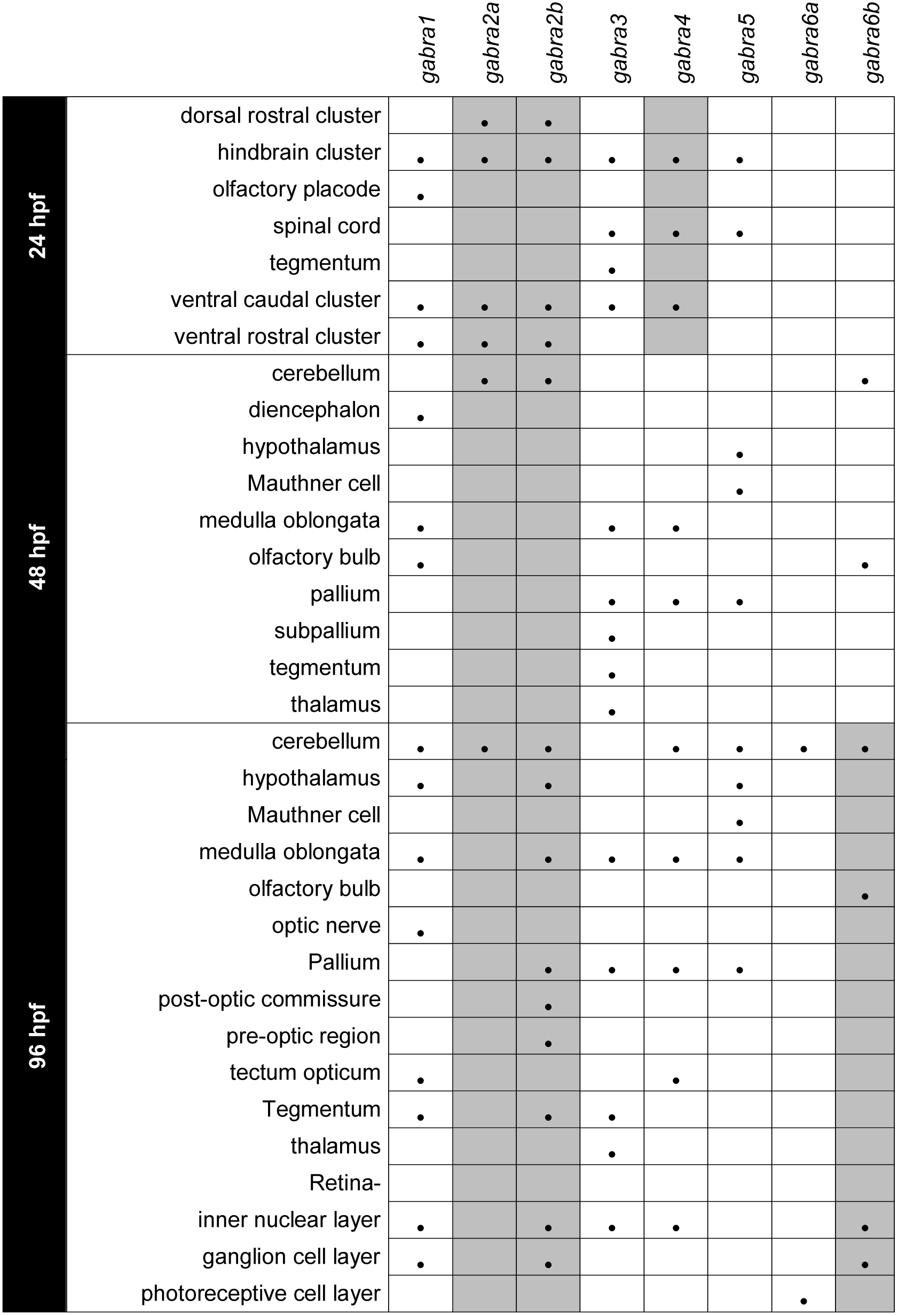
Expression of GABA_A_ receptor α subunits at 24, 48, and 96 hours post fertilization.

**Table 1** Shaded boxes indicate broad, diffuse expression was observed and dots indicate punctate expression was detected.

The expression patterns of GABA_A_ α subunits are well-conserved across species. The expression of five α subunits has been described in *Xenopus laevis* during development [48]. Consistent with our observations in zebrafish, α2 was found to be expressed broadly and early in development. In fact, α2 was found to be deposited in eggs as a maternal message. α6 showed the latest onset of expression during development and demonstrated robust photoreceptor expression. Likewise the expression patterns of α1, α3, and α5 are similar between frogs and zebrafish. These similarities in expression extend to mammalian systems. For example, in rat, similar to zebrafish α2, transcripts were detected relatively early in development and were widespread, and α6 is expressed much later in development and exhibits restricted expression [7]. One notable difference is that in mammals α6 transcripts were detected in the cerebellum and retinal expression has not been reported, whereas in zebrafish α6a exhibits robust retinal expression and α6b is more widespread. It is unclear how much these differences in expression between zebrafish and mammals can be attributed to the various development stages and tissues selected for investigation versus authentic differences in patterns of expression.

Developing zebrafish are a powerful system for neural circuit analysis and, more recently, they have been cultivated as a model of epilepsy [17, 49–53]. Although GABA_A_ receptors are robust regulators of many neural circuits and play a central role in epilepsy syndromes, GABA_A_ receptors have not been well characterized in zebrafish. By reporting the diversity of zebrafish GABA_A_ receptor subunits and describing the expression patterns of the α subfamily during early development, we have laid a foundation to leverage the strengths of the zebrafish system to investigate isoform specific roles of GABA_A_ receptors and further develop zebrafish as a model of epilepsy. In mammalian systems, genetic inactivation of individual α subunits has revealed surprisingly subtle defects compared to what would be predicted from pharmacological blockade. Compensatory upregulation of other subunits is thought to mask some of the effects of inactivating a single subunit [54, 55]. The ability easily mutate multiple genes in zebrafish may help overcome such compensatory changes in expression to shed new light into the subtype specific roles of GABA_A_ receptors.

## ACKNOWLEDGEMENTS

The authors thank members of the Downes lab, Kelly Anne McKeown at Westfield State University, and Josef Trapani at Amherst College for helpful advice and discussion.

## Supporting Information

**S1 Fig. RT-PCR results suggest that zebrafish contain only one α3 gene** (A) Schematic of a *gabra3* cDNA. The portions that are identical to *gabra3*-like and *gabra3* sequences are shaded green. The white region that connects the two was identified through RT-PCR. The location of the three primers used for PCR are shown as numbered arrows. (B) RT-PCR results. Primers 2 and 3 served as a positive control since they amplify sequence from the known *gabra3* region. Primers 1 and 3 amplify a previously unknown region that links *gabra* 3-like and *gabra3* sequence, showing that they are contained within same transcript and likely portions of the same gene.

**S1 Table. GABA_A_ α subunit antisense RNA probe information.** Note- Sizes are shown in base pairs. The probe start site is defined here as the start codon of the open reading frame. Negative values indicate that the probe contains 5’ untranslated sequence. Some probes were generated via RT-PCR with a primer that contains a RNA polymerase promoter and subcloning into vectors wasn’t performed.

## REFERENCES

1. Bowery NG, Smart TG. GABA and glycine as neurotransmitters: a brief history. Br J Pharmacol. 2006;147 Suppl 1:S109–19. Epub 2006/01/13. doi: 10.1038/sj.bjp.0706443. PubMed PMID: 16402094; PubMed Central PMCID: PMCPMC1760744.

2. Mohler H. GABA(A) receptor diversity and pharmacology. Cell Tissue Res. 2006;326(2):505–16. Epub 2006/08/29. doi: 10.1007/s00441-006-0284-3. PubMed PMID: 16937111.

3. Sigel E, Steinmann ME. Structure, function, and modulation of GABA(A) receptors. J Biol Chem. 2012;287(48):40224–31. Epub 2012/10/06. doi: 10.1074/jbc.R112.386664. PubMed PMID: 23038269; PubMed Central PMCID: PMCPMC3504738.

4. Chua HC, Chebib M. GABAA Receptors and the Diversity in their Structure and Pharmacology. Adv Pharmacol. 2017;79:1–34. Epub 2017/05/23. doi: 10.1016/bs.apha.2017.03.003. PubMed PMID: 28528665.

5. Olsen RW, Sieghart W. International Union of Pharmacology. LXX. Subtypes of gamma-aminobutyric acid(A) receptors: classification on the basis of subunit composition, pharmacology, and function. Update. Pharmacol Rev. 2008;60(3):243–60. Epub 2008/09/16. doi: 10.1124/pr.108.00505. PubMed PMID: 18790874; PubMed Central PMCID: PMCPMC2847512.

6. Laurie DJ, Seeburg PH, Wisden W. The distribution of 13 GABAA receptor subunit mRNAs in the rat brain. II. Olfactory bulb and cerebellum. J Neurosci. 1992;12(3):1063–76. Epub 1992/03/01. PubMed PMID: 1312132.

7. Laurie DJ, Wisden W, Seeburg PH. The distribution of thirteen GABAA receptor subunit mRNAs in the rat brain. III. Embryonic and postnatal development. J Neurosci. 1992;12(11):4151–72. PubMed PMID: 1331359.

8. Wisden W, Laurie DJ, Monyer H, Seeburg PH. The distribution of 13 GABAA receptor subunit mRNAs in the rat brain. I. Telencephalon, diencephalon, mesencephalon. J Neurosci. 1992;12(3):1040–62. Epub 1992/03/01. PubMed PMID: 1312131.

9. Delgado-Lezama R, Loeza-Alcocer E, Andres C, Aguilar J, Guertin PA, Felix R. Extrasynaptic GABA(A) receptors in the brainstem and spinal cord: structure and function. Curr Pharm Des. 2013;19(24):4485–97. Epub 2013/01/31. PubMed PMID: 23360278.

10. Simon J, Wakimoto H, Fujita N, Lalande M, Barnard EA. Analysis of the set of GABA(A) receptor genes in the human genome. J Biol Chem. 2004;279(40):41422–35. Epub 2004/07/20. doi: 10.1074/jbc.M401354200. PubMed PMID: 15258161.

11. Olsen RW, Sieghart W. GABA A receptors: subtypes provide diversity of function and pharmacology. Neuropharmacology. 2009;56(1):141–8. Epub 2008/09/02. doi: 10.1016/j.neuropharm.2008.07.045. PubMed PMID: 18760291; PubMed Central PMCID: PMCPMC3525320.

12. Tretter V, Moss SJ. GABA(A) Receptor Dynamics and Constructing GABAergic Synapses. Front Mol Neurosci. 2008;1:7. Epub 2008/10/24. doi: 10.3389/neuro.02.007.2008. PubMed PMID: 18946540; PubMed Central PMCID: PMCPMC2526003.

13. Fritschy JM, Panzanelli P. GABAA receptors and plasticity of inhibitory neurotransmission in the central nervous system. Eur J Neurosci. 2014;39(11):1845–65. Epub 2014/03/19. doi: 10.1111/ejn.12534. PubMed PMID: 24628861.

14. Farrant M, Nusser Z. Variations on an inhibitory theme: phasic and tonic activation of GABA(A) receptors. Nat Rev Neurosci. 2005;6(3):215–29. Epub 2005/03/02. doi: 10.1038/nrn1625. PubMed PMID: 15738957.

15. Egawa K, Fukuda A. Pathophysiological power of improper tonic GABA(A) conductances in mature and immature models. Front Neural Circuits. 2013;7:170. Epub 2013/10/30. doi: 10.3389/fncir.2013.00170. PubMed PMID: 24167475; PubMed Central PMCID: PMCPMC3807051.

16. Wolman M, Granato M. Behavioral genetics in larval zebrafish: learning from the young. Dev Neurobiol. 2012;72(3):366–72. Epub 2012/02/14. doi: 10.1002/dneu.20872. PubMed PMID: 22328273.

17. McLean DL, Fetcho JR. Movement, technology and discovery in the zebrafish. Curr Opin Neurobiol. 2011;21(1):110–5. Epub 2010/10/26. doi: 10.1016/j.conb.2010.09.011. PubMed PMID: 20970321; PubMed Central PMCID: PMCPMC3057273.

18. Baraban SC, Taylor MR, Castro PA, Baier H. Pentylenetetrazole induced changes in zebrafish behavior, neural activity and c-fos expression. Neuroscience. 2005;131(3):759–68. Epub 2005/02/26. doi: 10.1016/j.neuroscience.2004.11.031. PubMed PMID: 15730879.

19. Stehr CM, Linbo TL, Incardona JP, Scholz NL. The developmental neurotoxicity of fipronil: notochord degeneration and locomotor defects in zebrafish embryos and larvae. Toxicol Sci. 2006;92(1):270–8. Epub 2006/04/14. doi: 10.1093/toxsci/kfj185. PubMed PMID: 16611622.

20. Connaughton VP, Nelson R, Bender AM. Electrophysiological evidence of GABAA and GABAC receptors on zebrafish retinal bipolar cells. Vis Neurosci. 2008;25(2):139–53. Epub 2008/04/30. doi: 10.1017/S0952523808080322. PubMed PMID: 18442437; PubMed Central PMCID: PMCPMC4272819.

21. Del Bene F, Wyart C, Robles E, Tran A, Looger L, Scott EK, et al. Filtering of visual information in the tectum by an identified neural circuit. Science. 2010;330(6004):669–73. Epub 2010/10/30. doi: 10.1126/science.1192949. PubMed PMID: 21030657; PubMed Central PMCID: PMCPMC3243732.

22. Mack-Bucher JA, Li J, Friedrich RW. Early functional development of interneurons in the zebrafish olfactory bulb. Eur J Neurosci. 2007;25(2):460–70. Epub 2007/02/08. doi: 10.1111/j.1460-9568.2006.05290.x. PubMed PMID: 17284187.

23. Hubbard JM, Bohm UL, Prendergast A, Tseng PB, Newman M, Stokes C, et al. Intraspinal Sensory Neurons Provide Powerful Inhibition to Motor Circuits Ensuring Postural Control during Locomotion. Curr Biol. 2016;26(21):2841–53. Epub 2016/10/11. doi: 10.1016/j.cub.2016.08.026. PubMed PMID: 27720623.

24. Triller A, Rostaing P, Korn H, Legendre P. Morphofunctional evidence for mature synaptic contacts on the Mauthner cell of 52-hour-old zebrafish larvae. Neuroscience. 1997;80(1):133–45. Epub 1997/09/01. PubMed PMID: 9252227.

25. Roy B, Ali DW. Multiple types of GABAA responses identified from zebrafish Mauthner cells. Neuroreport. 2014;25(15):1232–6. Epub 2014/08/28. doi: 10.1097/WNR.0000000000000258. PubMed PMID: 25162782.

26. Cocco A, Ronnberg AM, Jin Z, Andre GI, Vossen LE, Bhandage AK, et al. Characterization of the gamma-aminobutyric acid signaling system in the zebrafish (Danio rerio Hamilton) central nervous system by reverse transcription-quantitative polymerase chain reaction. Neuroscience. 2017;343:300–21. Epub 2016/07/28. doi: 10.1016/j.neuroscience.2016.07.018. PubMed PMID: 27453477.

27. Hortopan GA, Dinday MT, Baraban SC. Spontaneous seizures and altered gene expression in GABA signaling pathways in a mind bomb mutant zebrafish. J Neurosci. 2010;30(41):13718–28. Epub 2010/10/15. doi: 10.1523/JNEUR0SCI.1887-10.2010. PubMed PMID: 20943912; PubMed Central PMCID: PMCPMC2962868.

28. Gonzalez-Nunez V. Role of gabra2, GABAA receptor alpha-2 subunit, in CNS development. Biochem Biophys Reports. 2015;3:11.

29. Baxendale S, Holdsworth CJ, Meza Santoscoy PL, Harrison MR, Fox J, Parkin CA, et al. Identification of compounds with anti-convulsant properties in a zebrafish model of epileptic seizures. Dis Model Mech. 2012;5(6):773–84. Epub 2012/06/26. doi: 10.1242/dmm.010090. PubMed PMID: 22730455; PubMed Central PMCID: PMCPMC3484860.

30. Thisse B, Thisse, C. Fast Release Clones: A High Throughput Expression Analysis. ZFIN Direct Data Submission. 2004.

31. Thompson JD, Higgins DG, Gibson TJ. CLUSTAL W: improving the sensitivity of progressive multiple sequence alignment through sequence weighting, position-specific gap penalties and weight matrix choice. Nucleic Acids Res. 1994;22(22):4673–80. Epub 1994/11/11. PubMed PMID: 7984417; PubMed Central PMCID: PMCPMC308517.

32. Guindon S, Dufayard JF, Lefort V, Anisimova M, Hordijk W, Gascuel O. New algorithms and methods to estimate maximum-likelihood phylogenies: assessing the performance of PhyML 3.0. Syst Biol. 2010;59(3):307–21. Epub 2010/06/09. doi: 10.1093/sysbio/syq010. PubMed PMID: 20525638.

33. Kimmel CB, Ballard WW, Kimmel SR, Ullmann B, Schilling TF. Stages of embryonic development of the zebrafish. Dev Dyn. 1995;203(3):253–310. PubMed PMID: 8589427.

34. Thisse C, Thisse B. High-resolution in situ hybridization to whole-mount zebrafish embryos. Nat Protoc. 2008;3(1):59–69. PubMed PMID: 18193022.

35. Mueller T, Wullimann MF. Atlas of early zebrafish brain development: a tool for molecular neurogenetics. Second edition. ed. xiii, 244 pages p.

36. Folgueira M, Bayley P, Navratilova P, Becker TS, Wilson SW, Clarke JD. Morphogenesis underlying the development of the everted teleost telencephalon. Neural Dev. 2012;7:32. Epub 2012/09/20. doi: 10.1186/1749-8104-7-32. PubMed PMID: 22989074; PubMed Central PMCID: PMCPMC3520737.

37. Sabaliauskas NA, Foutz CA, Mest JR, Budgeon LR, Sidor AT, Gershenson JA, et al. High-throughput zebrafish histology. Methods. 2006;39(3):246–54. Epub 2006/07/28. doi: 10.1016/j.ymeth.2006.03.001. PubMed PMID: 16870470.

38. Amores A, Force A, Yan YL, Joly L, Amemiya C, Fritz A, et al. Zebrafish hox clusters and vertebrate genome evolution. Science. 1998;282(5394):1711–4. Epub 1998/11/30. PubMed PMID: 9831563.

39. Postlethwait JH, Yan YL, Gates MA, Horne S, Amores A, Brownlie A, et al. Vertebrate genome evolution and the zebrafish gene map. Nat Genet. 1998;18(4):345–9. Epub 1998/04/16. doi: 10.1038/ng0498-345. PubMed PMID: 9537416.

40. Saint-Amant L, Drapeau P. Time course of the development of motor behaviors in the zebrafish embryo. J Neurobiol. 1998;37(4):622–32. PubMed PMID: 9858263.

41. Tsang SY, Ng SK, Xu Z, Xue H. The evolution of GABAA receptor-like genes. Mol Biol Evol. 2007;24(2):599–610. Epub 2006/12/01. doi: 10.1093/molbev/msl188. PubMed PMID: 17135332.

42. Darlison MG, Pahal I, Thode C. Consequences of the evolution of the GABA(A) receptor gene family. Cell Mol Neurobiol. 2005;25(3–4):607–24. Epub 2005/08/03. doi: 10.1007/s10571-005-4004-4. PubMed PMID: 16075381.

43. Martyniuk CJ, Aris-Brosou S, Drouin G, Cahn J, Trudeau VL. Early evolution of ionotropic GABA receptors and selective regimes acting on the mammalian-specific theta and epsilon subunits. PLoS One. 2007;2(9):e894. Epub 2007/09/20. doi: 10.1371/journal.pone.0000894. PubMed PMID: 17878929; PubMed Central PMCID: PMCPMC1975676.

44. Sinkkonen ST, Hanna MC, Kirkness EF, Korpi ER. GABA(A) receptor epsilon and theta subunits display unusual structural variation between species and are enriched in the rat locus ceruleus. J Neurosci. 2000;20(10):3588–95. Epub 2000/05/11. PubMed PMID: 10804200.

45. Higashijima S, Mandel G, Fetcho JR. Distribution of prospective glutamatergic, glycinergic, and GABAergic neurons in embryonic and larval zebrafish. J Comp Neurol. 2004;480(1):1–18. Epub 2004/10/30. doi: 10.1002/cne.20278. PubMed PMID: 15515020.

46. Kinkhabwala A, Riley M, Koyama M, Monen J, Satou C, Kimura Y, et al. A structural and functional ground plan for neurons in the hindbrain of zebrafish. Proc Natl Acad Sci U S A. 2011;108(3):1164–9. Epub 2011/01/05. doi: 10.1073/pnas.1012185108. PubMed PMID: 21199947; PubMed Central PMCID: PMCPMC3024665.

47. Downes GB, Waterbury JA, Granato M. Rapid in vivo labeling of identified zebrafish neurons. Genesis. 2002;34(3):196–202. PubMed PMID: 12395384.

48. Kaeser GE, Rabe BA, Saha MS. Cloning and characterization of GABAA alpha subunits and GABAB subunits in Xenopus laevis during development. Dev Dyn. 2011;240(4):862–73. Epub 2011/03/09. doi: 10.1002/dvdy.22580. PubMed PMID: 21384470; PubMed Central PMCID: PMCPMC3071254.

49. Fetcho JR, Higashijima S, McLean DL. Zebrafish and motor control over the last decade. Brain Res Rev. 2008;57(1):86–93. Epub 2007/09/11. doi: 10.1016/j.brainresrev.2007.06.018. PubMed PMID: 17825423; PubMed Central PMCID: PMCPMC2237884.

50. Friedrich RW, Genoud C, Wanner AA. Analyzing the structure and function of neuronal circuits in zebrafish. Front Neural Circuits. 2013;7:71. Epub 2013/05/01. doi: 10.3389/fncir.2013.00071. PubMed PMID: 23630467; PubMed Central PMCID: PMCPMC3632777.

51. Hale ME, Katz HR, Peek MY, Fremont RT. Neural circuits that drive startle behavior, with a focus on the Mauthner cells and spiral fiber neurons of fishes. J Neurogenet. 2016;30(2):89–100. Epub 2016/06/16. doi: 10.1080/01677063.2016.1182526. PubMed PMID: 27302612.

52. Griffin A, Krasniak C, Baraban SC. Advancing epilepsy treatment through personalized genetic zebrafish models. Prog Brain Res. 2016;226:195–207. Epub 2016/06/22. doi: 10.1016/bs.pbr.2016.03.012. PubMed PMID: 27323944.

53. Cunliffe VT. Building a zebrafish toolkit for investigating the pathobiology of epilepsy and identifying new treatments for epileptic seizures. J Neurosci Methods. 2016;260:91–5. Epub 2015/07/30. doi: 10.1016/j.jneumeth.2015.07.015. PubMed PMID: 26219659.

54. Kralic JE, Sidler C, Parpan F, Homanics GE, Morrow AL, Fritschy JM. Compensatory alteration of inhibitory synaptic circuits in cerebellum and thalamus of gamma-aminobutyric acid type A receptor alpha1 subunit knockout mice. J Comp Neurol. 2006;495(4):408–21. Epub 2006/02/18. doi: 10.1002/cne.20866. PubMed PMID: 16485284.

55. Peng Z, Zhang N, Chandra D, Homanics GE, Olsen RW, Houser CR. Altered localization of the delta subunit of the GABAA receptor in the thalamus of alpha4 subunit knockout mice. Neurochem Res. 2014;39(6):1104–17. Epub 2013/12/20. doi: 10.1007/s11064-013-1202-1. PubMed PMID: 24352815; PubMed Central PMCID: PMCPMC4024081.

